# Leveraging integrative toxicogenomic approach towards development of stressor-centric adverse outcome pathway networks for plastic additives

**DOI:** 10.1101/2024.03.27.586984

**Authors:** Ajaya Kumar Sahoo, Nikhil Chivukula, Shreyes Rajan Madgaonkar, Kundhanathan Ramesh, Shambanagouda Rudragouda Marigoudar, Krishna Venkatarama Sharma, Areejit Samal

## Abstract

Plastics are widespread pollutants found in atmospheric, terrestrial and aquatic ecosystems due to their extensive usage and environmental persistence. Plastic additives, that are utilized to achieve specific functionality in plastics, leach into the environment upon plastic degradation and pose considerable risk to ecological and human health. Limited knowledge concerning the presence of plastic additives throughout the plastic life cycle has hindered their effective regulation, thereby posing risks to product safety. In this study, we leveraged the adverse outcome pathway (AOP) framework to understand the mechanisms underlying plastic additives-induced toxicities. We first identified an exhaustive list of 6470 plastic additives from chemicals documented to be found in plastics. Next, we leveraged heterogenous toxicogenomics and biological endpoints data from five exposome-relevant resources, and identified associations between 1287 plastic additives and 322 complete and high quality AOPs within AOP-Wiki. Based on these plastic additive-AOP associations, we constructed a stressor-centric AOP network, wherein the stressors are categorized into 10 priority use sectors and AOPs are linked to 27 disease categories. We visualized the plastic additives-AOP network for each of the 1287 plastic additives and made them available in a dedicated website: https://cb.imsc.res.in/saopadditives/. Finally, we showed the utility of the constructed plastic additives-AOP network by identifying 28 highly relevant AOPs associated with benzo[a]pyrene, and thereafter, explored the associated toxicity pathways leading to respiratory and gastrointestinal system diseases in humans and developmental disorders in aquatic species. Overall, the constructed plastic additives-AOP network will enable regulatory risk assessment of plastic additives, thereby contributing towards a toxic-free circular economy for plastics.

## 1. Introduction

Plastics are the most widely produced synthetic chemicals, roughly constituting about 10% of solid waste generated globally (Geyer et al., 2017; Thompson et al., 2009). Extensive usage followed by improper waste management of plastics have made them ubiquitous pollutants in atmosphere, terrestrial and aquatic environments (Li et al., 2021; Parthasarathy et al., 2019). Plastics constitute (up to 50% by weight) various chemicals termed as plastic additives which aid in achieving its specific desirable properties (Hahladakis et al., 2018; Maes et al., 2023; UNEP, 2023). These plastic additives are not covalently bonded to plastic, and thus can be easily released into the environment throughout the plastic life cycle (Hermabessiere et al., 2017; Maddela et al., 2023; UNEP, 2023). Environmental exposure to such plastic additives has been observed to elicit various adverse health effects such as cancer, developmental defects, and endocrine disruptions in humans and other species alike (Maddela et al., 2023; Meeker et al., 2009; Oehlmann et al., 2008), but the lack of information on their presence throughout the plastics life cycle hampered their risk assessment and eventually the product safety (UNEP, 2023). Therefore, it is imperative to identify these plastic additives and perform their risk assessment to achieve a toxic-free circular economy for plastics.

In an attempt to improve and accelerate chemical toxicity testing in 21^st^ century, the US National Research Council in their report titled ‘Toxicity Testing in the 21st century: a vision and strategy’ advocated the use of new approach methodologies like *in vitro* and *in silico* high-throughput screening strategies (National Research Council, 2007). In accordance with the report’s suggestions, Ankley *et al*. (Ankley et al., 2010) proposed the conceptual toxicological framework namely, adverse outcome pathway (AOP), that sequentially organizes the stressor-induced biological response as Key Events (KEs) across various levels of biological organizations, thereby capturing the mechanism of stressor-induced toxicity (Ankley et al., 2010; Villeneuve et al., 2014a). In an AOP, the originating KE corresponds to the stressor-induced molecular event and is termed as molecular initiating event (MIE), whereas the terminal KE corresponds to the stressor-induced adverse effect and is termed as adverse outcome (AO) (Ankley et al., 2010; OECD, 2017; Villeneuve et al., 2014a). The sequential information in an AOP is represented in a linear setup where the flow of information is captured by directional links connecting different KEs (including MIEs and AOs), which are termed as Key Event Relationships (KERs) (Villeneuve et al., 2014a, 2014b; Vinken et al., 2017). Therefore, exploring toxicities induced by plastic additives using an AOP framework can streamline their risk assessment.

AOP-Wiki (https://aopwiki.org/) is the largest open-source repository which catalogues AOPs that are developed globally. In AOP-Wiki, an AOP is developed in a stressor-agnostic manner, where it captures a single toxicity pathway by linking a molecular perturbation (MIE) to an adverse effect (AO) of regulatory relevance (Knapen et al., 2018; Villeneuve et al., 2018). Thus, linking a stressor to different AOPs within AOP-Wiki can help identify all possible toxicity pathways associated with that stressor (Knapen et al., 2018).

Recently, we had leveraged heterogenous biological data from five exposome-relevant resources namely, ToxCast, Comparative Toxicogenomics Database (CTD) (Davis et al., 2023), DEDuCT (Karthikeyan et al., 2021, 2019), NeurotoxKb (Ravichandran et al., 2021b), and AOP-Wiki to extensively explore various toxicity pathways associated with inorganic cadmium-induced toxicity (Sahoo et al., 2024). Previously, Aguayo-Orozco *et al*. (Aguayo-Orozco et al., 2019) had utilized biological endpoint data of chemicals screened through several high throughout toxicity assays in ToxCast (Dix et al., 2007) to construct stressor-AOP network linking these chemicals to several developed AOPs within AOP-Wiki. Such a construction enabled exploration of the adverse effects associated with this chemical space from a mechanistic perspective (Aguayo-Orozco et al., 2019). Therefore, a data integrative approach can help identify AOPs within AOP-Wiki that are relevant for plastic additives-induced toxicity. Such plastic additive-AOP associations can aid in the development of stressor-centric AOP network for plastic additives that will provide a holistic view of plastic additives-induced adverse effects, thereby aiding in its regulatory decision-making.

In this study, we relied on the United Nations (UN) report titled ‘Chemicals in Plastics – A Technical Report’ (UNEP, 2023) that catalogued chemicals found in plastics from two independent studies by Aurisano *et al*. (Aurisano et al., 2021) and Wiesinger *et al*. (Wiesinger et al., 2021) and identified plastic additives among them based on their reported functions. First, we relied on AOP-Wiki, and systematically curated complete and connected high quality AOPs. Next, we leveraged the toxicogenomics and biological endpoints data from ToxCast, CTD, DEDuCT, NeurotoxKb, and AOP-Wiki to identify KEs associated with the curated plastic additives. Thereafter, based on these associated KEs, we constructed stressor-centric AOP network for plastic additives (designated as plastic additives-AOP network). Further, we linked the plastic additives to their corresponding priority use sectors and the AOPs to their corresponding disease categories in the constructed plastic additives-AOP network. Finally, we showed the utility of the constructed plastic additives-AOP network by identifying highly relevant plastic additive associated AOPs and leveraged published experimental evidence to explore plastic additive-induced toxicities including ecotoxicity. In sum, this is the first study leveraging heterogenous biological data to construct a large-scale stressor-centric AOP network for plastic additives.

## 2. Methods

### 2.1. Compilation and curation of plastic additives

Recently, the United Nations Environment Program (UNEP) published a report titled ‘Chemicals in Plastics – A Technical Report’ (UNEP, 2023) that provides an annex cataloguing over 13000 chemicals found in plastics and plastic manufacturing processes that were systematically curated by Aurisano *et al*. (Aurisano et al., 2021) and Wiesinger *et al*. (Wiesinger et al., 2021). Among the various chemicals in plastics, the compounds termed as plastic additives define the desirable properties in the final plastic product (UNEP, 2023). Plastic additives can constitute anywhere between 4%-50% by weight in plastics (UNEP, 2023), and can easily leach into the environment as they are not covalently bonded to the plastic polymers, thus posing a risk to human health and environment (Maddela et al., 2023). Here, we relied on the annex provided by the UNEP report to identify the different plastic additives (Figure 1).

**Figure 1:**
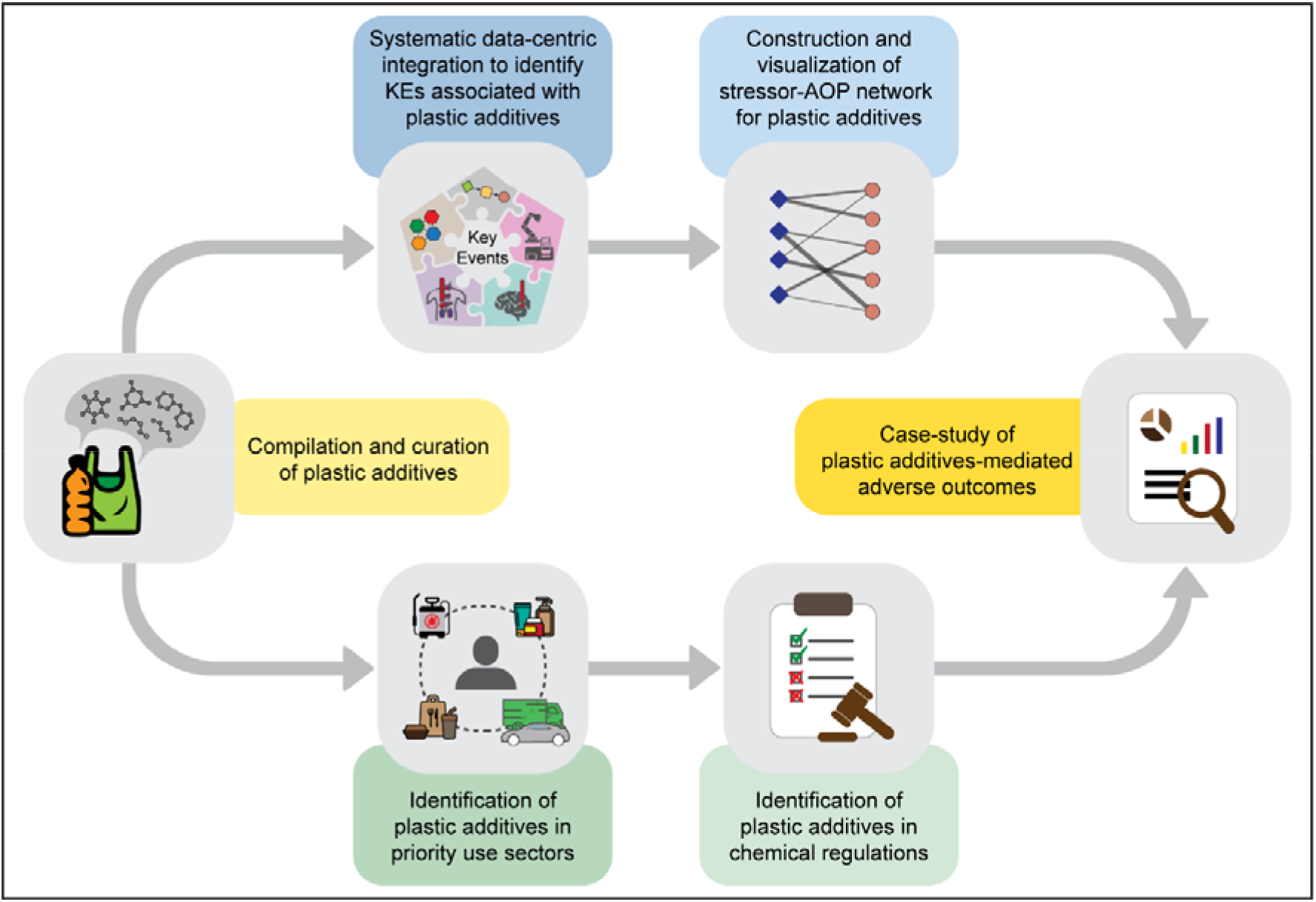
Summary of the workflow followed to identify plastic additives from chemicals found in plastics, followed by the exploration of their toxicity pathways through the construction of stressor-centric adverse outcome pathway (AOP) networks.

We first compiled the chemicals and their corresponding chemical abstracts service (CAS) registry numbers provided by the UNEP report. Thereafter, we relied on the CAS common chemistry web portal (https://commonchemistry.cas.org/) to identify synonymous CAS registry numbers and mapped them to their latest identifiers to remove redundancy and duplications in chemical identifiers listed in the UNEP report. In case the CAS identifier provided by the UNEP report is not present in the portal, we used the identifier provided by the UNEP report, and finally compiled a list of unique chemicals.

Next, we observed that various terms were used in the UNEP report to identify functions of different chemicals in plastics. Therefore, we standardized the vocabulary of the associated functions by relying on various sources (Al-Malaika et al., 2017; Coleman, 2017; Geyer et al., 2017; Hahladakis et al., 2018; Koleske et al., 2011; Pritchard, 2012; UNEP, 2023), and consequently identified different plastic additives among the compiled list of chemicals in plastics in the UNEP report (Supplementary Table S1). Through this extensive manual effort, we finally curated a list of 6470 plastic additives (Supplementary Table S2) from the chemicals compiled in the UNEP report, and leveraged them for further analysis.

### 2.2. Compilation of AOPs within AOP-Wiki

The AOP-Wiki (https://aopwiki.org/) is the largest publicly accessible repository, hosted by the Society for the Advancement of Adverse Outcome Pathways (SAAOP), which compiles and organizes various AOPs developed globally. An AOP is a linear framework that captures information on biological targets (called Molecular Initiating Events or MIEs) perturbed due to the action of an external stressor, and the cascade of events (called Key Events or KEs) following this perturbation that culminates into Adverse Outcomes (AOs) (Ankley et al., 2010; OECD, 2017, 2016). AOP-Wiki hosts this information on several AOPs documented in the form of KEs and the relationship between these KEs, known as Key Event Relationships (KERs), both of which are supported by scientific evidence.

In order to access the latest information available within AOP-Wiki, we downloaded the XML file (released on 1 January 2024) from ‘Project Downloads’ page in AOP-Wiki. Then, we utilized an in-house python script to parse the XML file and extract various information associated with AOPs like AOP identifier, AOP title, associated KEs (including MIEs and AOs) and KERs, linked stressors, status according to OECD and SAAOP, and biological applicability information such as taxonomy, sex and life-stage of the organism, and their corresponding weight of evidence. Additionally, we also extracted information associated with KEs like KE title, KE identifier, level of biological organization, action name, object name, object identifiers and process name, and information associated with KERs like upstream/downstream KEs, evidence for biological plausibility of KER, adjacency, and the extent of quantitative understanding of KER.

### 2.3. Identification of ‘high confidence AOPs’ within AOP-Wiki

Within AOP-Wiki, the AOPs are continuously updated based on current understanding and availability of novel experimental data, and thus, the AOPs are living documents (https://aopwiki.org/handbooks/4). Therefore, we relied on a systematic workflow developed in our previous work (Sahoo et al., 2024), to filter high quality and complete AOPs within AOP-Wiki (Supplementary Figure S1). First, we filtered out AOPs with SAAOP status as ‘archived’. Next, we manually checked and removed AOPs that have KE title as ‘unknown’ or lacked any KEs or KERs (Supplementary Figure S1). Next, we checked for the presence of disconnected components in AOPs using NetworkX library (Hagberg et al., 2008) in python, and manually updated and filtered out AOPs that contained disconnected components. Lastly, we checked the remaining AOPs for presence of MIEs, AOs and a directed path between MIE and AO, and filtered out AOPs that did not contain any such path (Supplementary Figure S1). This combined computational and manual effort led to the identification of 328 complete, connected and high quality AOPs within AOP-Wiki (last accessed on 15 February 2024) which we designate as ‘high confidence AOPs’ (Supplementary Figure S1; Supplementary Table S3). The 328 high confidence AOPs comprise 1107 unique KEs (Supplementary Table S4) and 1717 unique KERs (Supplementary Table S5).

### 2.4. Identification of KEs associated with plastic additives

Based on our previous work (Sahoo et al., 2024), we relied on a systematic and comprehensive data-centric integration method to identify the KEs within AOP-Wiki that are associated with plastic additives by utilizing toxicogenomics and biological endpoints data from five exposome-relevant resources: ToxCast (Dix et al., 2007), Comparative Toxicogenomics Database (CTD) (Davis et al., 2023) (https://ctdbase.org/), DEDuCT (Karthikeyan et al., 2021, 2019) (https://cb.imsc.res.in/deduct/), NeurotoxKb (Ravichandran et al., 2021b) (https://cb.imsc.res.in/neurotoxkb/) and AOP-Wiki (https://aopwiki.org/).

#### 2.4.1. Using ToxCast

US EPA’s ToxCast program provides high throughput *in vitro* bioactivity assay data for thousands of chemicals tested across several assays (Dix et al., 2007). Importantly, the ToxCast data includes assay annotations and information on associated bioprocess and genes that can aid in the identification of KEs (specifically MIEs) associated with the corresponding active chemical (Aguayo-Orozco et al., 2019; Knapen et al., 2018; Sahoo et al., 2024). First, we downloaded the latest ToxCast invitrodb version 4.1 (EPA, 2023) dataset from the US EPA repository (https://www.epa.gov/chemical-research/exploring-toxcast-data). Next, we retrieved chemicals and their corresponding assay endpoints from the ‘mc5-6_winning_model_fits-flags_invitrodb_v4_1_SEPT2023.csv’ file and filtered chemicals with active assay endpoints (‘hitc’ ≥ 0.9) (Feshuk et al., 2023). Furthermore, we retrieved the ‘activatory’ or ‘inhibitory’ response of these active chemicals by relying on the ‘top’ value of the corresponding winning model from the ‘mc4_all_model_fits_invitrodb_v4_1_SEPT2023.csv’ file (Feshuk et al., 2023).

Sometimes, chemicals exhibit their activity in a narrow range of concentrations that coincides with that of cell stress and cytotoxicity, thereby leading to non-specific activation of reporter genes. Such phenomena are termed as ‘cytotoxicity-associated bursts’ and can lead to inaccurate assay endpoint readings (Judson et al., 2016). Therefore, in this study, we identified such cytotoxicity-associated bursts for plastic additives tested within ToxCast, and did not consider those endpoints for mapping with KEs within AOP-Wiki (Supplementary Figure S2).

To identify cytotoxicity-associated bursts within ToxCast, Judson *et al*. (Judson et al., 2016) proposed the following Z-score metric:

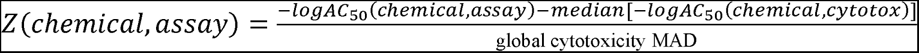

wherein, ‘logAC_50_(chemical,assay)’ is the logarithm of the AC_50_ value of the chemical in the assay, ‘logAC_50_(chemical,cytotox)’ is the logarithm of the AC_50_ value of the chemical in the corresponding cytotoxicity assay and the ‘global cytotoxicity MAD’ is the median of the MAD (median average deviations) of the logAC_50_(chemical,cytotox) distributions across all chemicals (Judson et al., 2016). For a given chemical, assays having Z-score values lying between +3 and -3 were considered as cytotoxicity-associated bursts (Judson et al., 2016). Here, we relied on this Z-score metric by Judson *et al*. to identify cytotoxicity-associated bursts corresponding to the plastic additives tested within ToxCast.

The ‘cytotox_invitrodb_v4_1_SEPT2023.xlsx’ file in ToxCast invitrodb version 4.1 provides the global cytotoxicity MAD and logAC_50_(chemical,cytotox) (‘cytotox_median_log’) for chemicals across various cytotoxicity assays (Feshuk et al., 2023). We retrieved these values for the plastic additives tested within ToxCast, computed their Z-scores and discarded assays that had a Z-score value lying between +3 and -3. Through this process, we identified 1108 assay endpoints associated with 1327 plastic additives, and proceeded to map them to KEs within AOP-Wiki (Supplementary Figure S2).

First, we retrieved the genes associated with these 1108 assay endpoints from the ‘assay_annotations_invitrodb_v4_1_SEPT2023.xlsx’ file and the details of assay endpoint-gene mappings from ‘assay_gene_mappings_invitrodb_v4_1_SEPT2023’ file. Next, we leveraged KE-gene annotations provided by Saarimäki *et al*. (Saarimäki et al., 2023) to identify gene sets associated with KEs having biological level of organization as either molecular or cellular. Thereafter, we mapped these KEs to ToxCast assay endpoints based on gene overlaps, and manually filtered the mappings based on the assay endpoint descriptions. Through this extensive toxicogenomics based manual curation, we obtained 212 assay endpoints mapped to 115 KEs for 1129 of the 6470 curated plastic additives (Supplementary Figure S2).

#### 2.4.2. Using CTD

Comparative Toxicogenomics Database (CTD) (Davis et al., 2023) (https://ctdbase.org/) is one of the largest toxicogenomics resources that compiles data on chemical-gene/protein, chemical-disease, and gene-disease associations from published literature. The concept of chemical (C), gene (G), phenotype (P) and disease (D) tetramers, i.e. CGPD-tetramers, was proposed to understand the phenotypes and diseases that result from the interaction of chemicals with genes (Davis et al., 2020). Based on our previous work (Sahoo et al., 2024), we retrieved the CGPD-tetramers associated with plastic additives within CTD, and leveraged them to identify the associated KEs within AOP-Wiki.

First, we downloaded the CTD’s January 2024 release and constructed the CGPD-tetramers for the plastic additives based on the workflow proposed in our previous work (Sahoo et al., 2024). This resulted in the identification of 124496 tetramers comprising 258 chemicals, 2932 genes, 1489 phenotypes and 690 diseases (Supplementary Table S6). Furthermore, we generated the immediate neighbor GO terms for the CGPD-tetramer phenotype GO terms using the GOSim package (Fröhlich et al., 2007) available in R programming language. Thereafter, we overlapped the GO terms with the process identifiers of KEs within AOP-Wiki, and manually inspected to identify 307 KEs associated with 266 phenotypes for 241 of the 6470 curated plastic additives (Supplementary Table S7). We also manually inspected the disease terms to identify 157 KEs associated with 315 diseases for 232 of the 6470 curated plastic additives (Supplementary Table S7).

#### 2.4.3. Using DEDuCT and NeurotoxKb

DEDuCT (Karthikeyan et al., 2021, 2019) (https://cb.imsc.res.in/deduct/) is one of the largest databases that compiles curated information on endocrine disrupting chemicals (EDCs) and their corresponding endocrine-mediated endpoints from published literature. Therefore, we compiled the endocrine-mediated endpoints corresponding to plastic additives within DEDuCT, and considered them to find associated KEs within AOP-Wiki. We manually inspected the endpoints and titles of KEs within AOP-Wiki, and identified 165 KEs that are associated with 188 endocrine-mediated endpoints for 203 of the 6470 curated plastic additives (Supplementary Table S7).

NeurotoxKb (Ravichandran et al., 2021b) (https://cb.imsc.res.in/neurotoxkb/) is a manually curated resource on mammalian neurotoxicity associated endpoints of environmental chemicals curated from published literature. Therefore, we compiled the neurotoxic endpoints corresponding to plastic additives within NeurotoxKb, and considered them to find associated KEs within AOP-Wiki. We manually inspected the neurotoxic endpoints and KEs within AOP-Wiki, and identified 25 KEs that are associated with 24 neurotoxic endpoints for 92 of the 6470 curated plastic additives (Supplementary Table S7).

#### 2.4.4. Using AOP-Wiki

AOP-Wiki also catalogues the stressor information for each AOP, where there exists well documented evidence of such stressor(s) showing response at multiple KEs, including MIEs (https://aopwiki.org/handbooks/4). Therefore, we relied on the stressor information within AOP-Wiki to identify KEs associated with plastic additives. We retrieved information on stressors associated with each AOP, and identified 33 AOPs to be associated with 42 of the 6470 curated plastic additives (Supplementary Table S7). Thereafter, we identified 178 KEs in these 33 AOPs that are associated with plastic additives.

Overall, we identified 688 KEs that are associated with 1314 plastic additives (out of the 6470 plastic additives in our curated list) through the integration of heterogenous toxicogenomics and biological endpoints data from five exposome-relevant resources: ToxCast, CTD, DEDuCT, NeurotoxKb and AOP-Wiki (Supplementary Table S7).

### 2.5. Compilation of chemical lists for priority use sectors of plastic additives

Globally, plastics are used across different scales and for various applications. The UNEP report (UNEP, 2023) has identified 10 priority use sectors, based on the likelihood of exposure of chemicals in plastic products in these sectors to humans and environment. The 10 priority use sectors include ‘Toys and other children’s products’, ‘Furniture’, ‘Packaging including food contact materials’, ‘Electrical and electronic equipment’, ‘Transport’, ‘Personal care and household products’, ‘Medical devices’, ‘Building materials’, ‘Synthetic textiles’, and ‘Agriculture, aquaculture and fisheries’. To identify the plastic additives being used in each of the priority use sectors, we first compiled the list of chemicals in use in each of these sectors.

Chemical and Products Database (CPDat) (Dionisio et al., 2018) (https://www.epa.gov/chemical-research/chemical-and-products-database-cpdat) is among the largest resources that catalogues the presence of chemicals in various consumer products. For each product, CPDat assigns a Product Use Category (PUC) based on the general category and product type mentioned in the original data source (Dionisio et al., 2018). Here, we relied on CPDat to identify the chemicals in use in each of the 10 priority use sectors. We accessed the CPDat data file (Williams, 2021) (last accessed on 15 February 2024) to compile the list of chemicals associated with different PUCs, and identified 20 PUCs to be grouped under the different priority use sectors (Supplementary Figure S3; Supplementary Table S8).

The CompTox Chemicals Dashboard (Williams et al., 2021, 2017) (https://comptox.epa.gov/dashboard/) is one of the largest public repositories that provides access to different lists of chemicals associated with projects, publications, source databases or collections (https://comptox.epa.gov/dashboard/chemical-lists). Here, we queried the chemical lists based on their description, and identified chemical lists associated with the different priority use sectors (Supplementary Figure S3; Supplementary Table S8). Furthermore, we compiled chemicals from an in-house repository namely, Fragrance Chemicals in Children’s Products (FCCP) (Ravichandran et al., 2022a) (https://cb.imsc.res.in/fccp) as chemicals found in the use sector ‘Toys and other children’s products’. Finally, we compiled the chemicals in use in each of the priority use sectors, and identified plastic additives present in each sector (Supplementary Table S8).

### 2.6. Construction and visualization of the stressor-AOP network

Stressor-AOP network provides a holistic view of chemical perturbances across different AOPs (Aguayo-Orozco et al., 2019). To better understand the perturbances caused by the different plastic additives, we constructed a stressor-AOP network as a bipartite graph that linked various plastic additives to different AOPs within AOP-Wiki. In order to obtain high confidence associations between plastic additives and AOPs, we relied only on the curated list of 328 high confidence AOPs (Supplementary Table S3).

To obtain the stressor-AOP network for plastic additives, we initially linked plastic additives to AOPs if they share at least one associated KE, and thereafter, characterized each link between a stressor and an AOP based on the coverage score and level of relevance. Coverage score of a stressor-AOP link is defined as the ratio of number of KEs within that AOP associated with the stressor to the total number of KEs within that AOP (Chai et al., 2021). Coverage score is a real number that takes a value between 0 and 1 and we denote this score as the edge weight of linkage between a stressor and an AOP in our stressor-AOP network. Next, we realized that the plastic additives were associated with different AOPs with varying levels of relevance. Therefore, we propose the following five-level criterion to qualitatively understand the relevance of associations:

- *Level 1*: The stressor is associated with at least one KE within an AOP, where the KE is neither MIE nor AO within that AOP
- *Level 2*: The stressor is associated with at least one AO within an AOP, but not associated with any MIE within that AOP
- *Level 3*: The stressor is associated with at least one MIE within an AOP, but not associated with any AO within that AOP
- *Level 4*: The stressor is associated with at least one MIE and one AO within an AOP
- *Level 5*: The stressor is associated with at least one MIE and one AO within an AOP and there exists a directed path between the associated MIE and AO

Supplementary Table S9 contains all the data on the stressor-AOP network constructed for plastic additives, including the coverage scores and levels of relevance for each of the stressor-AOP links. We visualized this stressor-AOP network of plastic aditives using Cytoscape (Shannon et al., 2003).

## 3. Results and discussion

### 3.1. Exploration of the curated list of plastic additives

Plastic additives are chemicals that are added to plastics to achieve specific desirable properties in the end product (Hahladakis et al., 2018; Pritchard, 2012). External stress on such products can cause the separation of these additives, thereby leading to their release into the environment and eventually posing risks to humans and ecosystems (Maddela et al., 2023). In this study, we curated a list of plastic additives from chemicals found in plastics (Supplementary Table S2) and explored their potential risks by systematically integrating the associated heterogenous biological endpoints in the context of adverse outcome pathway (AOP) framework (Figure 1).

We relied on the chemicals provided in the UN report ‘Chemicals in Plastics – A Technical Report’ and identified 6470 chemicals as plastic additives based on their functions (Methods; Supplementary Table S2). We observed that many plastic additives (3217 of 6470) provide a variety of functions to the plastics, with colorants being the most frequently associated function (3675 of 6470) (Methods; Supplementary Table S2). Further, we observed that majority of plastic additives (4309 of 6470) are found in products made by different priority use sectors, of which 3963 additives are found in the use sector ‘Packaging, including food contact materials’ (Methods; Supplementary Table S2).

Next, we relied on the United States high production volume (USHPV) chemical list (https://comptox.epa.gov/dashboard/chemical-lists/EPAHPV) and Organisation for Economic Co-operation and Development high production volume (OECD HPV) chemical list (https://hpvchemicals.oecd.org/ui/Search.aspx) and identified 2084 of 6470 plastic additives to be high production volume (HPV) chemicals (Supplementary Table S2). Notably, among these HPV plastic additives, we found 154 additives to be known endocrine disrupting chemicals (EDCs) with experimental evidence for endocrine disruption in humans or rodents from DEDuCT (Karthikeyan et al., 2021, 2019), and 101 additives as potential carcinogens based on IARC monographs on identification of carcinogenic hazards to humans (https://monographs.iarc.who.int/list-of-classifications/) (Supplementary Table S2). Furthermore, we observed that 215 additives are identified as substances of very high concern (SVHC) (https://echa.europa.eu/candidate-list-table) by European Chemical Agency and 412 additives are prohibited for use as per REACH regulation (Supplementary Table S2). Figure 2a shows the distribution of HPV, SVHC and REACH prohibited plastic additives across the 10 priority use sectors.

**Figure 2:**
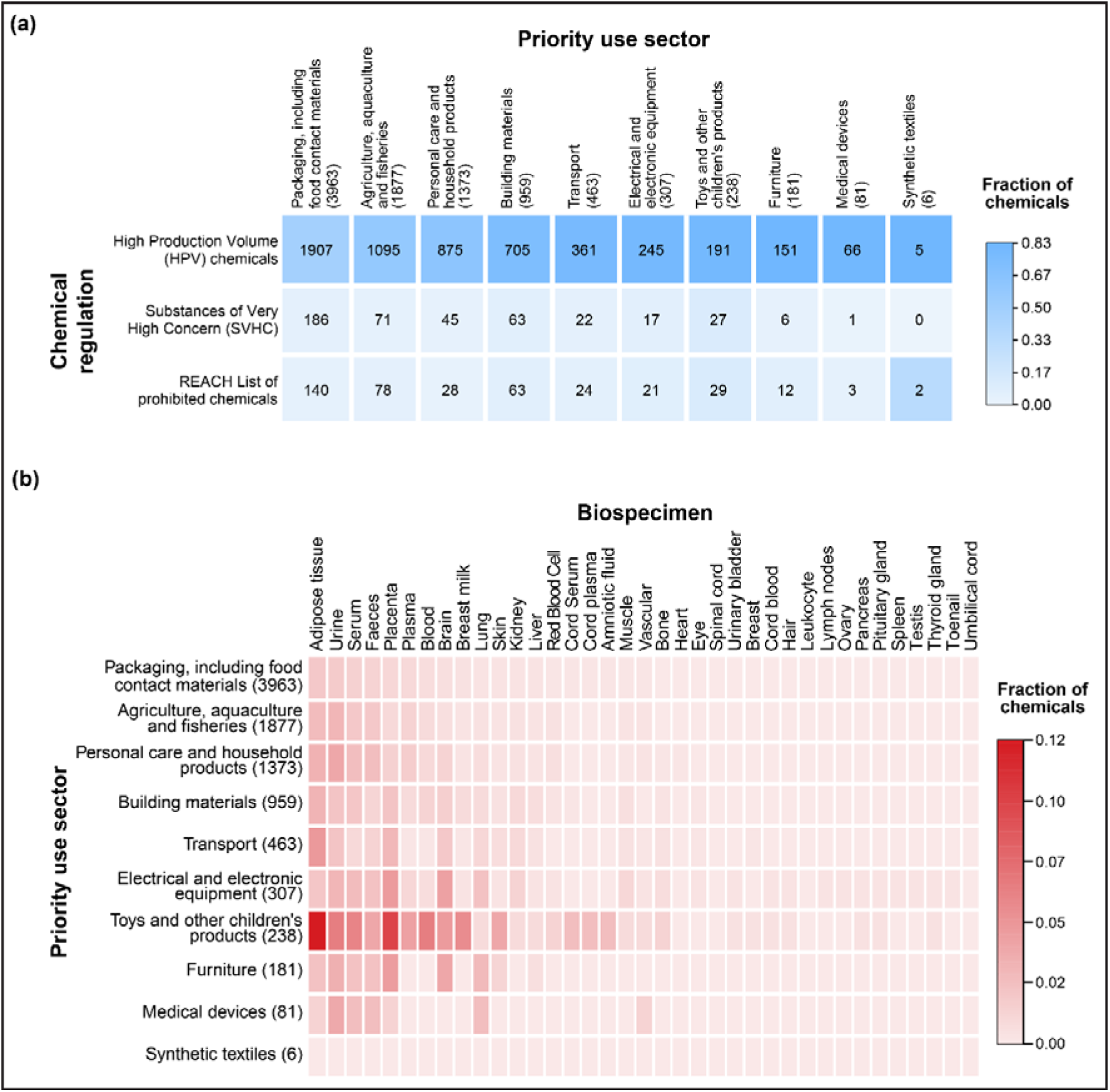
Identification of plastic additives in different chemical regulations and human biospecimens. **(a)** Heatmap depicting the presence of plastic additives from 10 priority use sectors in chemical regulations. The number of the plastic additives from the priority use sector in each of the chemical regulations is denoted in the heatmap. **(b)** Heatmap depicting the presence of plastic additives from 10 priority use sectors in different human biospecimens based on published exposure studies.

### 3.2. Plastic additives are accumulated in various human biospecimens

Humans are exposed to various plastic additives via direct contact, inhalation or ingestion, which can eventually accumulate in different human tissues and potentially lead to various adverse health effects (Meeker et al., 2009; UNEP, 2023). In order to explore the plastic additives detected in various human biospecimens, we relied on two databases namely, Tissue-specific Exposome Atlas (TExAs) (Ravichandran et al., 2021a) (https://cb.imsc.res.in/texas/) and Exposome-Explorer (Neveu et al., 2020) (http://exposome-explorer.iarc.fr/) which have compiled the presence of environmental chemicals as xenobiotics in different human tissues from published exposure studies. Although, these two databases have compiled information from limited human exposure studies, they have documented 204 of the 6470 plastic additives to be accumulated as xenobiotics in 37 different human biospecimens (Supplementary Table S2; Figure 2b). Moreover, we observed that plastic additives from 9 of the 10 priority use sectors have been documented as xenobiotics in human biospecimens namely, faeces, serum, urine, lung, placenta, and adipose tissue (Figure 2b). Note, the use sector ‘Synthetic textiles’ comprised the least number of additives (6 chemicals) in our curated list of 6470 additives, and there are no published exposure studies wherein their presence was detected in different human biospecimens.

### 3.3. Stressor-AOP network for plastic additives

Stressor-AOP networks provide a panoramic visualization of the different AOPs associated with stressors of interest, and help in understanding the stressor-induced adverse biological effects (Aguayo-Orozco et al., 2019). In this study, we therefore constructed stressor-AOP network to understand the various adverse effects induced by plastic additives. First, we followed a systematic approach that involved data-centric integration of heterogenous toxicogenomics and biological endpoints data from five exposome-relevant resources namely, ToxCast, CTD, DEDuCT, NeurotoxKb and AOP-Wiki, and identified 688 KEs within AOP-Wiki to be associated with 1314 of the 6470 plastic additives (Methods; Supplementary Table S7). Thereafter, we curated 328 high confidence AOPs within AOP-Wiki and mapped them to plastic additives if at least one KE within that AOP is associated with the plastic additive. Based on these plastic additive-AOP associations, we constructed a plastic additives-centric bipartite stressor-AOP network comprising of two types of nodes namely, 1287 plastic additives and 322 high confidence AOPs, and 46243 stressor-AOP links as edges between the two types of nodes, and we designate this bipartite network as plastic additives-AOP network. Notably, we observed that AOP-Wiki documented only 37 of the 1287 plastic additives in the constructed stressor-AOP network to be associated with 27 of the 322 high confidence AOPs in the network.

Next, we leveraged the KEs associated with plastic additives to compute the coverage score for the stressor-AOP links in the plastic additives-AOP network and observed that 20 plastic additives are associated with all the KEs (coverage score = 1.0) in 15 high confidence AOPs, and these stressor-AOP links were otherwise not documented in AOP-Wiki (Methods; Supplementary Table S9). Moreover, we calculated the levels of relevance for the stressor-AOP links in the plastic additives-AOP network and observed that 27189 links between 1155 plastic additives and 288 AOPs are classified as Level 1, 4236 links between 345 plastic additives and 241 AOPs are classified as Level 2, 14187 links between 1152 plastic additives and 139 AOPs are classified as Level 3, and 631 links between 118 plastic additives and 98 AOPs are classified as Level 5 (Methods; Supplementary Table S9). Note, the stressor-AOP links with Level 4 relevance were also satisfied by Level 5 criterion, and therefore, there are no stressor-AOP links with Level 4 relevance in the constructed network (Methods; Supplementary Table S9).

Next, we relied on the standardized disease ontology provided in Disease Ontology (Schriml et al., 2022) database (https://disease-ontology.org/do) to classify the AOPs based on their adverse outcomes (AOs). Based on the standardized ontology, we classified 322 AOPs into 26 disease classes based on their AOs (Supplementary Tables S9-S10). Note that 125 of the 322 AOPs could not be classified under any standardized ontology provided by Disease Ontology, and we therefore marked them as ‘unclassified’. Importantly, we observed that cancer is the most represented disease category comprising 40 of the 322 AOPs in the plastic additives-AOP network (Supplementary Table S9). Finally, we have linked the plastic additives to their corresponding priority use sectors and the AOPs to their corresponding disease categories in the plastic additives-AOP network (Supplementary Table S9).

We relied on the graph visualization software Cytoscape (Shannon et al., 2003) to visualize the plastic additives-AOP network for each of the 1287 plastic additives, and make them available on a dedicated website: https://cb.imsc.res.in/saopadditives/. In the website, the plastic additives are grouped based on their priority use sectors. For instance, the priority use sector ‘Toys and other children’s products’ consists of 162 plastic additives, 301 AOPs and 8300 stressor-AOP links, wherein 4696 links between 148 plastic additives and 265 AOPs are classified as Level 1, 1460 links between 75 plastic additives and 170 AOPs are classified as Level 2, 1928 links between 144 plastic additives 117 AOPs are classified as Level 3, and 216 links between 30 plastic additives and 58 AOPs are classified as Level 5. Figure 3 shows a portion of the plastic additives-AOP network, comprising Level 5 stressor-AOP links for plastic additives in the use sector ‘Toys and other children’s products’.

**Figure 3:**
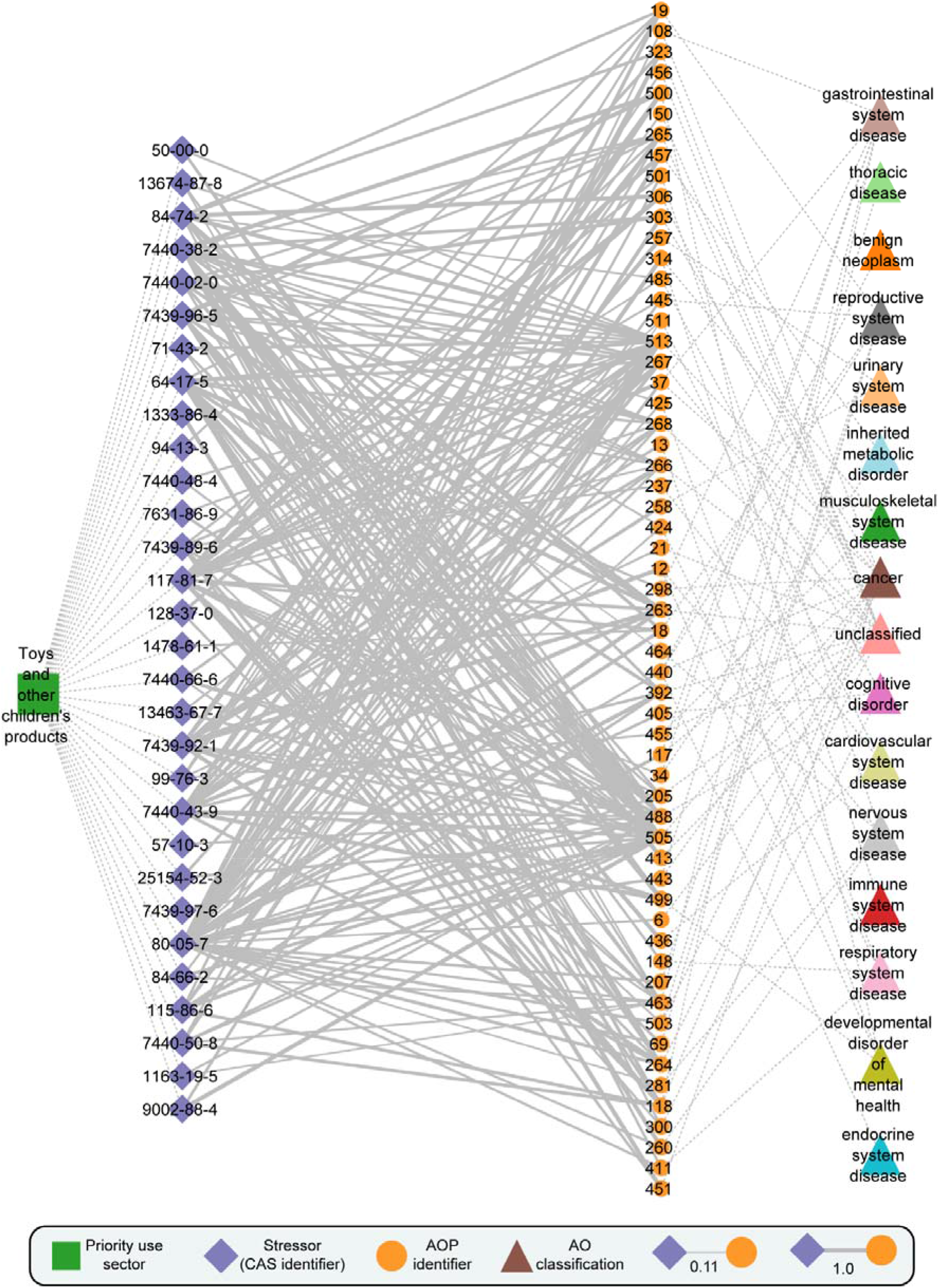
Visualization of a stressor-centric AOP network for plastic additives in the priority use sector ‘Toys and other children’s products’, along with the disease classifications of adverse outcomes (AOs) in AOPs. In the stressor-AOP network, only edges or stressor-AOP links with Level 5 relevance are shown. Further, the edges or stressor-AOP links are weighted based on their coverage score, i.e., the fraction of KEs within AOP that are linked with the plastic additives.

Additionally, the constructed plastic additives-AOP network can highlight the adverse outcomes induced by plastic additives across different use sectors. Figure 4 shows the disease categories linked with 10 priority use sectors for plastic additives in the plastic additives-AOP network with Level 5 relevance, where cancer is the most represented disease category.

**Figure 4:**
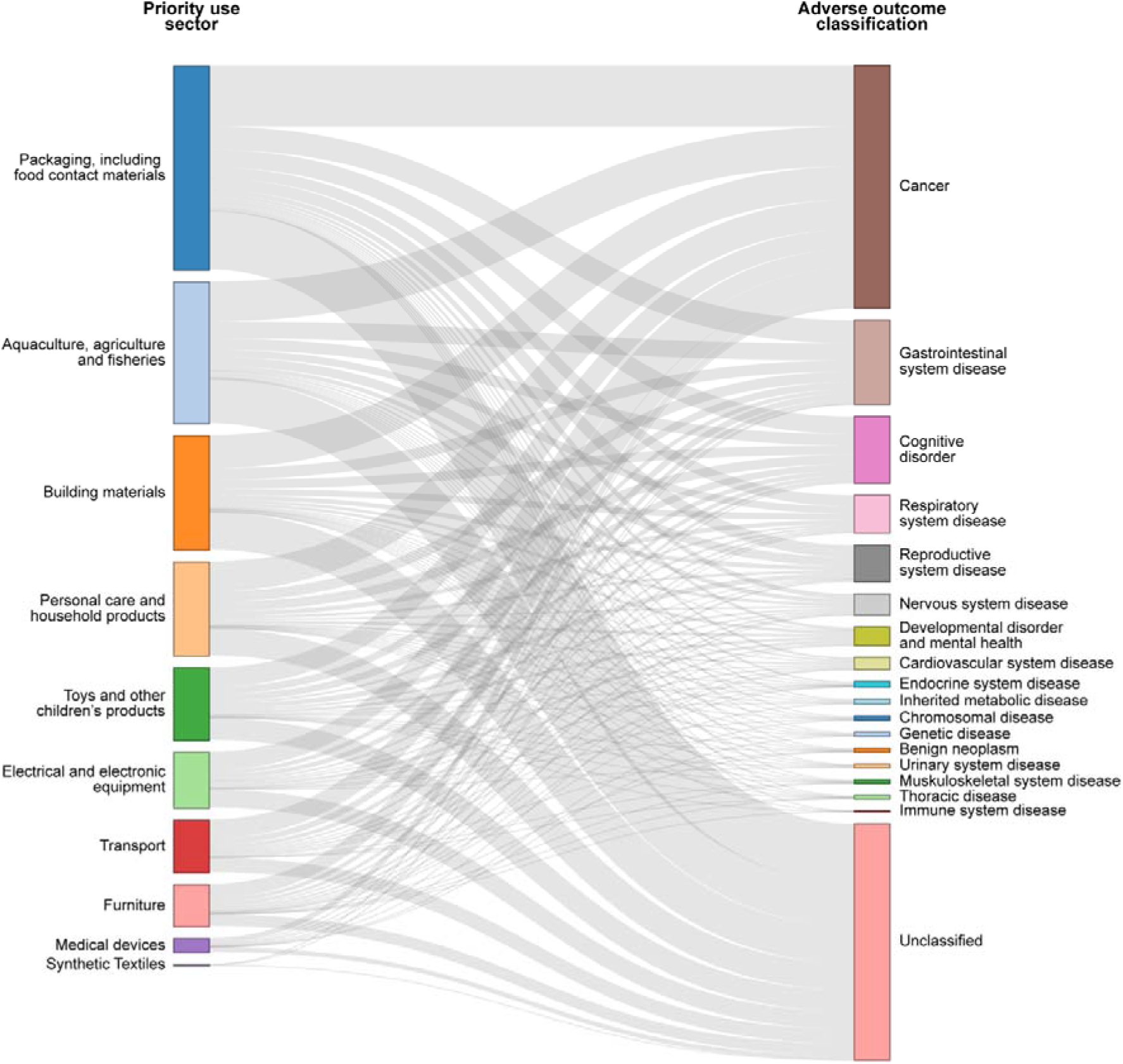
Aggregate visualization of the disease categories associated with AOPs in plastic additives stressor-AOP network with Level 5 relevance for each of the 10 priority use sectors.

### 3.4. Stressor-AOP network reveals highly relevant AOPs associated with plastic additives

A stressor-AOP network can help identify most relevant AOPs associated with each stressor which can further highlight the complexity and diversity among toxicity pathways induced by that stressor (Knapen et al., 2018). Here, we considered stressor-AOP links from the constructed plastic additives-AOP network with Level 5 relevance and coverage score threshold of 0.4, and identified 107 of the 1287 plastic additives to be associated with 88 of the 322 AOPs through 526 stressor-AOP links (Methods; Supplementary Table S9). Note the coverage score threshold of 0.4 denotes that at least 40% of the KEs in that AOPs are linked with the stressor (Chai et al., 2021; Sahoo et al., 2024). Notably, 15 of these 107 plastic additives are associated with more than 10 AOPs (Table 1). Among these 15 plastic additives, 14 are documented as EDCs in DEDuCT, and 10 are documented as carcinogens in IARC monographs (Supplementary Table S2).

**Table 1:** The top 15 plastic additives based on the associated number of highly relevant AOPs in the plastic additives-AOP network with Level 5 stressor-AOP links.

| **Plastic additive** | **Number of highly relevant stressor-AOP links** | **Number of stressor-AOP links not present in AOP-Wiki** |
| --- | --- | --- |
| Benzo[a]pyrene (CAS:50-32-8) | 28 | 28 |
| Bisphenol A (CAS:80-05-7) | 27 | 26 |
| Bis(2-ethylhexyl) phthalate (CAS:117-81-7) | 19 | 17 |
| Arsenic (CAS:7440-38-2) | 16 | 13 |
| Ethanol (CAS:64-17-5) | 15 | 13 |
| Perfluorooctanoic Acid (CAS:335-67-1) | 14 | 14 |
| Triclosan (CAS:3380-34-5) | 14 | 13 |
| Cadmium (CAS:7440-43-9) | 14 | 11 |
| Cadmium chloride (CAS:10108-64-2) | 13 | 12 |
| Lead (CAS:7439-92-1) | 13 | 10 |
| Acrylamide (CAS:79-06-1) | 13 | 13 |
| Pentachlorophenol (CAS:87-86-5) | 12 | 11 |
| Manganese (CAS:7439-96-5) | 11 | 9 |
| Oxygen (CAS:7782-44-7) | 11 | 11 |
| Lead acetate (CAS:301-04-2) | 11 | 11 |

Among the AOPs associated with these 15 plastic additives, we observed that majority of the AOPs are identified through our systematic data integrative approach. Notably, we observed that AOP:263, AOP:264, AOP:265, AOP:267, and AOP:268 are shared among all 15 plastic additives (Table 1). Moreover, we observed that these five AOPs share the same MIE ‘Decrease, Coupling of oxidative phosphorylation’ (KE:1446) and AO ‘Decrease, Growth’ (KE:1521), while AOP:263 titled ‘Uncoupling of oxidative phosphorylation leading to growth inhibition via decreased cell proliferation’ is endorsed by Working Group of the National Coordinators of the Test Guidelines Programme (WNT) and the Working Party on Hazard Assessment (WPHA) under the OECD AOP development programme.

An AOP network constructed from stressor-specific AOPs can highlight interactions among the associated AOPs, thereby aiding in the assessment of stressor-induced toxicity (Chai et al., 2021; Pogrmic-Majkic et al., 2022; Rugard et al., 2020; Sahoo et al., 2024; Yang et al., 2022). Among the 15 plastic additives, we observed that Benzo[a]pyrene (B[a]P or CAS:50-32-8) has the largest number of associated AOPs (28 AOPs), all of which were solely identified through our systematic data integrative approach (Table 1). These 28 AOPs are classified under various disease categories namely, cancer, gastrointestinal system disease, reproductive system disease, respiratory system disease, cognitive disorder, thoracic disease, and musculoskeletal system disease (Supplementary Table S9). Previously, Yang *et al*. (Yang et al., 2022) had constructed an AOP network for B[a]P-induced toxicity, but they relied only on CTD to identify KEs associated with B[a]P and focused only on B[a]P-induced male reproductive damages. Therefore, we relied on 28 AOPs associated with B[a]P-induced toxicity (which we designate as B[a]P-AOPs) and constructed an AOP network to explore various adverse effects associated with B[a]P.

Next, we computed the cumulative weight of evidence (WoE) for each of these 28 B[a]P-AOPs based on their KER information to assess the biological plausibility (Ravichandran et al., 2022b; Sahoo et al., 2024). We observed that 9 of these 28 B[a]P-AOPs have ‘High’ cumulative WoE and 6 B[a]P-AOPs have ‘Moderate’ cumulative WoE (Supplementary Table S11). Moreover, we observed that many of these 28 B[a]P-AOPs are applicable across various species and developmental stages (Supplementary Table S11). Figure 5 shows the undirected AOP network representation of the 28 B[a]P-AOPs, where nodes represent B[a]P-AOPs and the edges represent the existence of shared KEs between two AOPs. We observed that the B[a]P-AOPs form three connected components (with two or more AOPs) and two isolated nodes, where the largest connected component (labelled C1) comprises 18 B[a]P-AOPs. We considered the largest connected component (C1) for further analysis.

**Figure 5:**
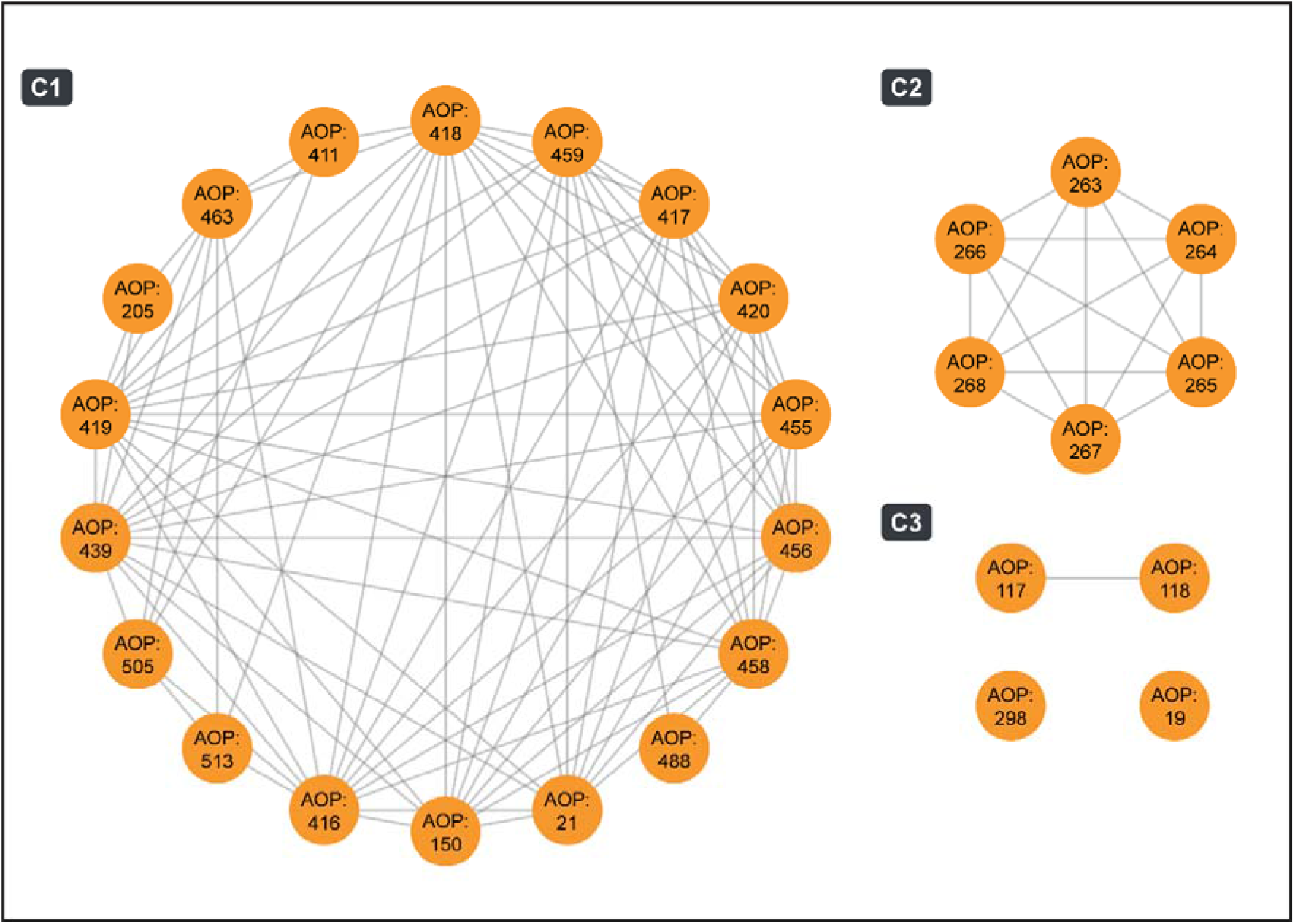
Undirected network of B[a]P-AOPs. Each node corresponds to B[a]P-AOP and an edge between two nodes denotes that the two AOPs share at least one KE. This undirected network has 3 connected components (with two or more nodes) which are labeled as C1, C2, and C3, and 2 isolated nodes.

### 3.5. Exploration of toxicity pathways in AOP network constructed from B[a]P-relevant AOPs

We constructed and visualized a directed AOP network to explore the interactions among the B[a]P-AOPs present in the largest connected component C1 (Figure 6). We observed that the directed network comprised 66 unique KEs (including 7 MIEs and 11 AOs) and 99 unique KERs (Figure 6; Supplementary Table S12). Among the 66 KEs, 36 KEs were associated with B[a]P-induced toxicity through our systematic data integrative approach, of which 5 are MIEs and 10 are AOs (Figure 6). Notably, we observed that the toxicity pathway originating from MIE ‘Activation, AhR’ (KE:18), passing through KEs ‘Altered gene expression, NRF2 dependent antioxidant pathway’ (KE:1917) and ‘Increase, Cell Proliferation’ (KE:870), and eventually terminating at AO ‘Lung Cancer’ (KE:1670) consists of 4 of the 36 KEs associated with B[a]P-induced toxicity (Figure 6). Upon further inspection, we identified that this toxicity pathway was captured in the AOP:420 titled ‘Aryl hydrocarbon receptor activation leading to lung cancer through sustained NRF2 toxicity pathway’, which was systematically built and supported by extensive literature survey and experimental data on B[a]P (Jin et al., 2022).

**Figure 6:**
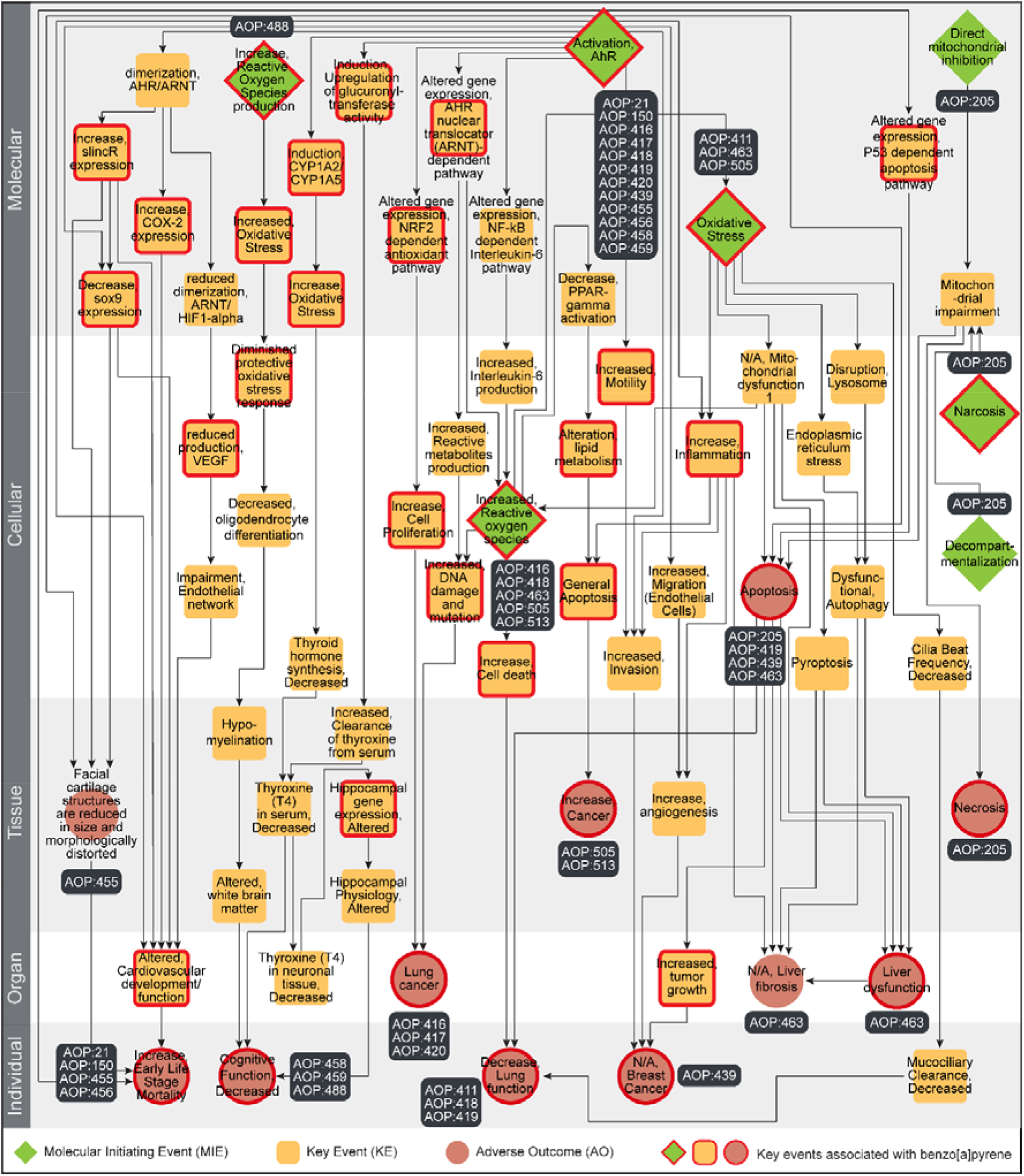
Directed network corresponding to the largest component in the undirected B[a]P-AOP network comprising 66 KEs and 99 KERs. Among the 66 KEs, 7 are categorized as MIEs (denoted as diamond), 11 are categorized as AOs (denoted as circle), and the remaining 48 are categorized as KEs (denoted as rounded square). The 36 KEs (including MIEs and AOs) associated with B[a]P are marked in ‘red’. In this figure, the 66 KEs are arranged vertically according to their level of biological organization.

Next, we computed different node-centric network measures to explore various features of this directed network. We observed that the MIE ‘Activation, AhR’ (KE:18) has the highest out-degree of 16, while the KE ‘Altered, Cardiovascular development/function’ (KE:317) and AO ‘N/A, Liver fibrosis’ (KE:344) have the highest in-degree of 5 (Supplementary Table S12). The MIE ‘Increased, Reactive oxygen species’ (KE:1115) has the highest betweenness centrality value, denoting that several toxicity pathways are passing through it in this network (Supplementary Figure S4) (Villeneuve et al., 2018). The KEs ‘Induction, CYP1A2/CYP1A5’ (KE:850) and ‘Altered gene expression, NF-kB dependent Interleukin-6 pathway’ (KE:1921), and MIEs ‘Activation, AhR’ (KE:18) and ‘Increase, Reactive Oxygen Species production’ (KE:257) have the highest eccentricity, denoting that they are the most remotely placed KEs in this network (Supplementary Figure S5) (Takes and Kosters, 2011).

Finally, we relied on artificial intelligence (AI) based tool, AOP-helpFinder (Jaylet et al., 2023; Jornod et al., 2022) (https://aop-helpfinder.u-paris-sciences.fr/), and Abstract Sifter (Baker et al., 2017) (https://comptox.epa.gov/dashboard/chemical/pubmed-abstract-sifter/) to screen published literature and manually identified novel associations between the B[a]P-induced toxicities and the remaining 30 KEs in the directed AOP network (Supplementary Table S13). Additionally, we compiled auxiliary evidence for the 36 KEs that were associated with B[a]P-induced toxicity through our systematic data integrative approach. Furthermore, we compiled information on the type of evidence and the reported toxicity dosage values of B[a]P exposure from these published evidence (Supplementary Table S13). To conclude, we performed two case studies to explore both the human-relevant and ecotoxicology-relevant B[a]P-induced toxicity pathways from this directed AOP network.

#### 3.5.1. Toxicity pathway linking B[a]P exposure to Liver fibrosis in humans

Liver fibrosis, which results from chronic damage to the liver, is a characteristic of many chronic liver diseases (Bataller and Brenner, 2005). Previously, exposome-based studies had found a significant association between environmental chemicals such as B[a]P and different liver diseases (Barouki et al., 2023; Cheung et al., 2020). Here, we observed an emergent B[a]P-induced toxicity pathway originating from MIE ‘Activation, AhR’ (KE:18) and terminating at AO ‘N/A, Liver fibrosis’ (KE:344). Therefore, we relied on this emergent toxicity pathway to understand the rationale behind B[a]P-induced liver fibrosis in humans.

Various *in vivo* and *in vitro* experiments in human cell lines and rodents had shown that B[a]P induces different downstream processes through the activation of AhR (Almendarez-Reyna et al., 2023; Jin et al., 2022; Lou et al., 2022; Tsai et al., 2015). Subsequently, B[a]P exposure has been observed to induce oxidative stress in cells through increased interleukin 6 (IL-6) production as a result of activated NF-κB signaling pathway (Jin et al., 2022; Malik et al., 2018; Zheng et al., 2022). The oxidative stress caused by B[a]P exposure has been studied as a cause for disruption of lysosomes, eventually leading to dysfunctional autophagy (Gorria et al., 2008; Li et al., 2023). Finally, it has been shown that B[a]P exposure can induce different fibrotic pathways, including dysfunctional autophagy, in human hepatic models (Hill and Wang, 2022; Yan et al., 2021). In conclusion, by leveraging various published evidence, we were able to explore a potential toxicity pathway that links B[a]P-induced toxicity with liver fibrosis in humans.

#### 3.5.2. Toxicity pathway linking B[a]P exposure to early life-stage mortality in aquatic organisms

B[a]P is found in large quantities in different aquatic environments due to various anthropogenic activities and waste discharges from both household and industries (Bukowska et al., 2022; Sathikumaran et al., 2022). B[a]P is a toxic pollutant, and drastically affects various aquatic organisms, including economically relevant fish (Nacci et al., 2002; Sathikumaran et al., 2022; Seemann et al., 2015). Here, we observed that the AOP titled ‘Aryl hydrocarbon receptor activation leading to early life stage mortality via sox9 repression induced impeded craniofacial development’ (AOP:455), with biological applicability for developmental effects in aquatic species, has been identified as a B[a]P-AOP (Supplementary Table S11). Moreover, this B[a]P-AOP is part of the largest connected component, and is currently included in the OECD work plan (Supplementary Table S1). Therefore, we relied on this AOP to understand the rationale behind B[a]P-induced ecotoxicological effects in aquatic organisms.

Independent *in vivo* experiments in zebrafish and clam have shown that B[a]P exposure alters gene expression patterns through activation of AhR and subsequent dimerization of AhR and ARNT in affected tissues (Bugiak and Weber, 2009; Wang et al., 2020). It has been shown that B[a]P exposure in zebrafish facilitates the recruitment of AhR-dependent long noncoding RNA (*slincR*) to *sox9b* 5’ UTR, eventually repressing its transcription (Garcia et al., 2018). *sox9b* is an important transcription factor involved in chondrocyte differentiation during zebrafish development (Dalcq et al., 2012). Subsequently it has been shown that B[a]P exposure induces alteration in expression patterns of genes involved in chondrogenesis, thereby leading to improper craniofacial skeleton development and eventually early life stage mortality in zebrafish (He et al., 2011; Seemann et al., 2015). In conclusion, by leveraging various published evidence, we were able to explore a potential toxicity pathway that links B[a]P-induced toxicity with early life stage mortality in aquatic species.

## 4. Conclusions

Plastic additives are chemicals that are easily released into the environment from various plastic products. The lack of information on their presence in various plastic products pose a challenge in evaluating their risks, thereby hindering proper regulatory measures. In this study, we constructed a stressor-centric AOP network of plastic additives to aid in their regulatory decision making. First, we relied on the UN report titled ‘Chemicals in Plastics – A Technical Report’ and identified 6470 plastic additives based on the reported chemical functions. Next, we systematically integrated heterogenous toxicogenomics and biological endpoints data from five exposome-relevant resources namely, ToxCast, CTD, DEDuCT, NeurotoxKb, and AOP-Wiki and identified 688 KEs within AOP-Wiki to be associated with 1314 plastic additives. Further, we systematically curated 328 high confidence AOPs from AOP-Wiki and linked them to plastic additives based on overlapping KE associations. Here, we identified 322 high confidence AOPs to be associated with 1287 plastic additives while AOP-Wiki only documented 37 of the 1287 plastic additives to be associated with 27 of the 322 high confidence AOPs. Next, we constructed the stressor-AOP network for plastic additives (designated as plastic additives-AOP network) with varying levels of associations, where the plastic additives are categorized into 10 priority use sectors and the AOPs are linked with 27 disease classes. We visualized the plastic additives-AOP network for each of the 1287 plastic additives and made them available in a dedicated website: https://cb.imsc.res.in/saopadditives/. Finally, we showed the utility of the constructed plastic additives-AOP network by identifying highly relevant AOPs (with Level 5 relevance and coverage score threshold of 0.4) associated with plastic additives. In particular, we identified 28 highly relevant AOPs associated with Benzo[a]pyrene (B[a]P) and relied on published experimental evidence to explore B[a]P-induced respiratory and gastrointestinal system diseases in humans and developmental disorders in aquatic species.

However, due to limited information on their presence in various use sectors, we were able to identify only 4309 of the 6470 plastic additives across 10 priority use sectors. Further, due to the paucity of plastic additive exposure studies, we were able to associate only 1287 of the 6470 plastic additives to AOPs within AOP-Wiki. We observed that 197 of the 322 high confidence AOPs (associated with the 1287 plastic additives) capture toxicity pathways leading to human relevant adverse effects. Moreover, the toxicogenomics approach followed in this study relied majorly on mammalian-centric biological data, thereby limiting its scope to explore various ecotoxicological events.

Nonetheless, we have constructed the first and most comprehensive stressor-AOP network for plastic additives that has enabled the exploration of different toxicity mechanisms underlying plastic additives-specific adverse outcomes. Further, we observed that many of these plastic additives are produced in high volumes globally and are documented to cause endocrine disruptions. Notably, we observed that these additives can accumulate in various human tissues as xenobiotics, suggesting that prolonged exposure to plastic additives can lead to highly deleterious effects in different organ systems (Maddela et al., 2023; Sorci and Loiseau, 2022). Importantly, the constructed plastic additives-AOP network was useful in the identification of highly relevant AOPs for plastic additives which highlighted plastic additives-induced emergent toxicity pathways. In sum, this is the first study that utilizes the AOP framework to explore the various adverse effects associated with plastic additives, enabling their risk assessment and contributing towards their regulatory decision-making.

## Data availability

The data associated with this study is contained in the article or in the supplementary material or in the associated website: https://cb.imsc.res.in/saopadditives/.

## Acknowledgements

The authors would like to thank K. Dhineka, M. Sambandam and Pravakar Mishra for discussions. Areejit Samal would like to acknowledge funding from the Department of Atomic Energy (DAE), Government of India via Apex project to The Institute of Mathematical Sciences (IMSc) Chennai. The funders have no role in study design, data collection, data analysis, manuscript preparation or decision to publish.

## CRediT author contribution statement

**Ajaya Kumar Sahoo:** Conceptualization, Data Compilation, Data Curation, Formal Analysis, Software, Visualization, Writing; **Nikhil Chivukula:** Conceptualization, Data Compilation, Data Curation, Formal Analysis, Software, Visualization, Writing; **Shreyes Rajan Madgaonkar:** Conceptualization, Data Compilation, Data Curation, Formal Analysis, Software, Visualization, Writing; **Kundhanathan Ramesh:** Data Compilation, Data Curation, Formal Analysis, Visualization, Writing; **Shambanagouda Rudragouda Marigoudar:** Formal Analysis, Writing; **Krishna Venkatarama Sharma:** Formal Analysis, Writing; **Areejit Samal:** Conceptualization, Supervision, Formal Analysis, Writing.

## Supplementary Tables

**Supplementary Table S1:** This table contains the list of standardized functions for plastic additives. For each function, the table provides the synonymous function names (separated by ‘|’ symbol) as given in the annex of the UN report (https://www.unep.org/resources/report/chemicals-plastics-technical-report), description of the function and the corresponding references (separated by ‘|’ symbol).

**Supplementary Table S2:** This table contains the list of 6470 plastic additives which are curated from the annex provided by the UN report (https://www.unep.org/resources/report/chemicals-plastics-technical-report). For each additive, the table provides the corresponding information on its primary CAS identifier, synonymous CAS identifiers (separated by ‘|’ symbol) as given in the annex, chemical name as given in the annex, standardized functions (separated by ‘|’ symbol), associated priority use sectors (separated by ‘|’ symbol), its presence as a high production volume (HPV) chemical as per US HPV or OECDHPV, its presence as substances of very high concern, its presence in the REACH list of prohibited chemicals, its presence as a potential carcinogen in IARC monographs on identification of carcinogenic hazards to humans, its presence as an EDC in DEDuCT, and its presence in human biospecimens (separated by ‘|’ symbol).

**Supplementary Table S3:** This table contains the curated list of 328 high confidence adverse outcome pathways (AOPs) from AOP-Wiki. For each AOP, the table provides the corresponding information on AOP identifier, AOP title, Organisation for Economic Co-operation and Development (OECD) status, and Society for the Advancement of Adverse Outcome Pathways (SAAOP) status.

**Supplementary Table S4:** This table contains information on the 1107 Key Events (KEs) present in the curated list of 328 high confidence AOPs. For each KE, the table provides the corresponding information on KE identifier, KE title, level of biological organization, and associated AOP identifier(s) (separated by ‘|’ symbol). Note that the KE identifiers starting with 10000 were manually assigned by the authors as the KE was mentioned in the AOP page but was not assigned an identifier in AOP-Wiki.

**Supplementary Table S5:** This table contains information on Key Event Relationships (KERs) present in each of the curated list of 328 high confidence AOPs. For each AOP, the table provides the AOP identifier, corresponding KER identifier, upstream KE identifier, downstream KE identifier, MIE(s) among upstream and downstream KEs (separated by ‘|’ symbol), AO(s) among upstream and downstream KEs (separated by ‘|’ symbol), adjacency of KER, weight of evidence of KER, and quantitative understanding of KER. Note that the KER identifiers starting with 10000 were manually assigned by the authors as the KER was mentioned in the AOP page but was not assigned an identifier in AOP-Wiki.

**Supplementary Table S6:** This table contains information on 124496 chemical-gene-phenotype-disease (CGPD) tetramers constructed for the plastic additives from Comparative Toxicogenomics Database (CTD). For each tetramer, the table provides the corresponding information on the CAS identifier for the chemical, NCBI gene identifier, NCBI gene name, phenotype identifier, phenotype name, MESH disease identifier, and MESH disease name.

**Supplementary Table S7:** This table contains information on 688 Key Events (KEs) that are associated with plastic additives. For each KE, the table provides corresponding information on KE identifier, CAS identifiers (separated by ‘|’) for associated additives, source(s) from which the associations are inferred (separated by ‘|’ symbol), CTD phenotype(s) (separated by ‘|’ symbol), CTD disease(s) (separated by ‘|’ symbol), ToxCast assay endpoint(s) (separated by ‘|’ symbol), endocrine-mediated endpoint(s) from DEDuCT (separated by ‘|’ symbol), and neurotoxic endpoint(s) from NeurotoxKb (separated by ‘|’ symbol).

**Supplementary Table S8:** This table contains information on the curated chemical category or list for each of the 10 priority use sectors. For each priority use sector, the table provides corresponding information on the chemical category or list name, their description, source, link, and number of chemicals. Further, the table provides the number of unique chemicals for each priority use sector and number of overlapped chemicals with the curated list of plastic additives.

**Supplementary Table S9:** This table provides the edge list for the complete stressor-AOP network for plastic additives. For each edge in the network, the table provides the CAS identifier for the stressor (plastic additive), plastic additive associated priority use sectors (separated by ‘|’ symbol), AOP identifier, classification of the adverse outcomes (AOs) in the AOP, and computed coverage score and level of relevance.

**Supplementary Table S10:** This table contains information on disease standardized ontology from the Disease Ontology database for the adverse outcomes (AOs) in AOPs. For each AO, the table provides the KE identifier, KE title, level of biological organization, disease names (separated by ‘|’ symbol) in Disease Ontology database, disease identifiers (separated by ‘|’ symbol) in Disease Ontology database and disease classes (separated by ‘|’ symbol) in Disease Ontology database. Note, if an AO could not be classified from Disease Ontology database, then we marked it as ‘unclassified’.

**Supplementary Table S11:** This table contains information on biological applicability and associated evidence for each of the 28 B[a]P-AOPs. For each AOP, the table provides the corresponding AOP identifier, computed cumulative weight of evidence (WoE), life-stage applicability of AOP [‘High’, ‘Moderate’, ‘Low’, ‘Not Specified’] (separated by ‘|’ symbol), taxonomic applicability of AOP [‘High’, ‘Moderate’, ‘Low’, ‘Not Specified’] (separated by ‘|’ symbol), and sex applicability of AOP [’High’, ‘Moderate’, ‘Low’, ‘Not Specified’] (separated by ‘|’ symbol).

**Supplementary Table S12:** This table contains information on the computed network statistics for the 66 Key Events (KEs) present in the largest connected component (C1) of the B[a]P-AOP network. For each KE, the table provides the corresponding KE identifier, KE title, KE type, in-degree value, out-degree value, eccentricity value, betweenness centrality value (rounded up to 4 decimal values), and convergence information.

**Supplementary Table S13:** This table contains information on the literature evidence of the association with B[a]P for each of the 66 KEs present in the largest connected component (C1) of the B[a]P-AOP network. For each KE, the table provides the corresponding KE identifier, KE title, level of biological organization, KE type, organism in which B[a]P exposure is studied, study type, dosage information, abbreviated description of association with B[a]P, and the corresponding PubMed identifier of the literature evidence.

## Supplementary Figures

**Supplementary Figure S1:** Workflow to filter high confidence adverse outcome pathways (AOPs) from AOP-Wiki by employing computation and manual curation in conjunction. (Adapted from Sahoo *et al*., 2024)

**Supplementary Figure S2:** Workflow to identify KEs from AOP-Wiki which are mapped to the active assay endpoints of plastic additives within ToxCast.

**Supplementary Figure S3:** Mapping of chemical or category lists from CompTox Chemicals Dashboard and Chemical and Products Database (CPDat) with the 10 priority use sectors of plastic additives.

**Supplementary Figure S4:** Directed network corresponding to the largest connected component (C1) in the B[a]P-AOP network, where the KEs (including MIEs and AOs) are colored based on their betweenness centrality values. The 36 KEs (including MIEs and AOs) associated with B[a]P are marked in ‘red’. In this figure, the 66 KEs are arranged vertically according to their level of biological organization.

**Supplementary Figure S5:** Directed network corresponding to the largest connected component (C1) in the B[a]P-AOP network, where the KEs (including MIEs and AOs) are colored based on their eccentricity values. The 36 KEs (including MIEs and AOs) associated with B[a]P are marked in ‘red’. In this figure, the 66 KEs are arranged vertically according to their level of biological organization.

