## Supplementary Figure for "Leveraging integrative toxicogenomic approach towards development of stressor-centric adverse outcome pathway networks for plastic additives"

### Supplementary Figures S1-S5

### for

### Leveraging integrative toxicogenomic approach towards development of stressor-centric adverse outcome pathway networks for plastic additives

Ajaya Kumar Sahoo[a,b,1], Nikhil Chivukula[a,b,1], Shreyes Rajan Madgaonkar[a,b], Kundhanathan Ramesh[a], Shambanagouda Rudragouda Marigoudar[c], Krishna Venkatarama Sharma[c], Areejit Samal[a,b,*]

[a] *The Institute of Mathematical Sciences (IMSc), Chennai, India*

[b] *Homi Bhabha National Institute (HBNI), Mumbai, India*

[c] *National Centre for Coastal Research, Ministry of Earth Sciences, Government of India, Pallikaranai, Chennai, India*

[1]A.K.S. and N.C. contributed equally to this work and should be considered as Joint-First authors

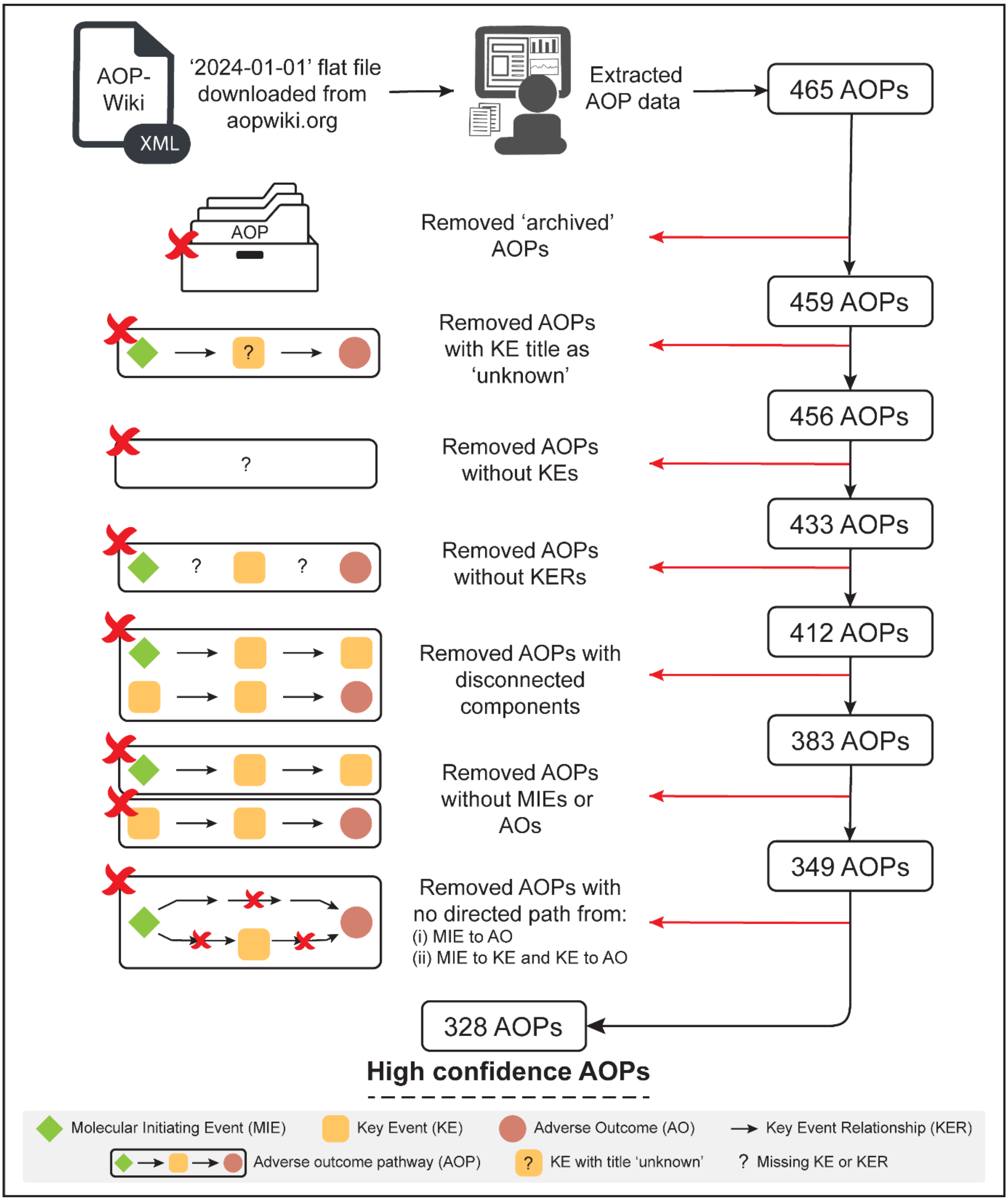


**Figure S1:** Workflow to filter high confidence adverse outcome pathways (AOPs) from AOP-Wiki by employing computation and manual curation in conjunction. (Adapted from Sahoo *et al*., 2024)

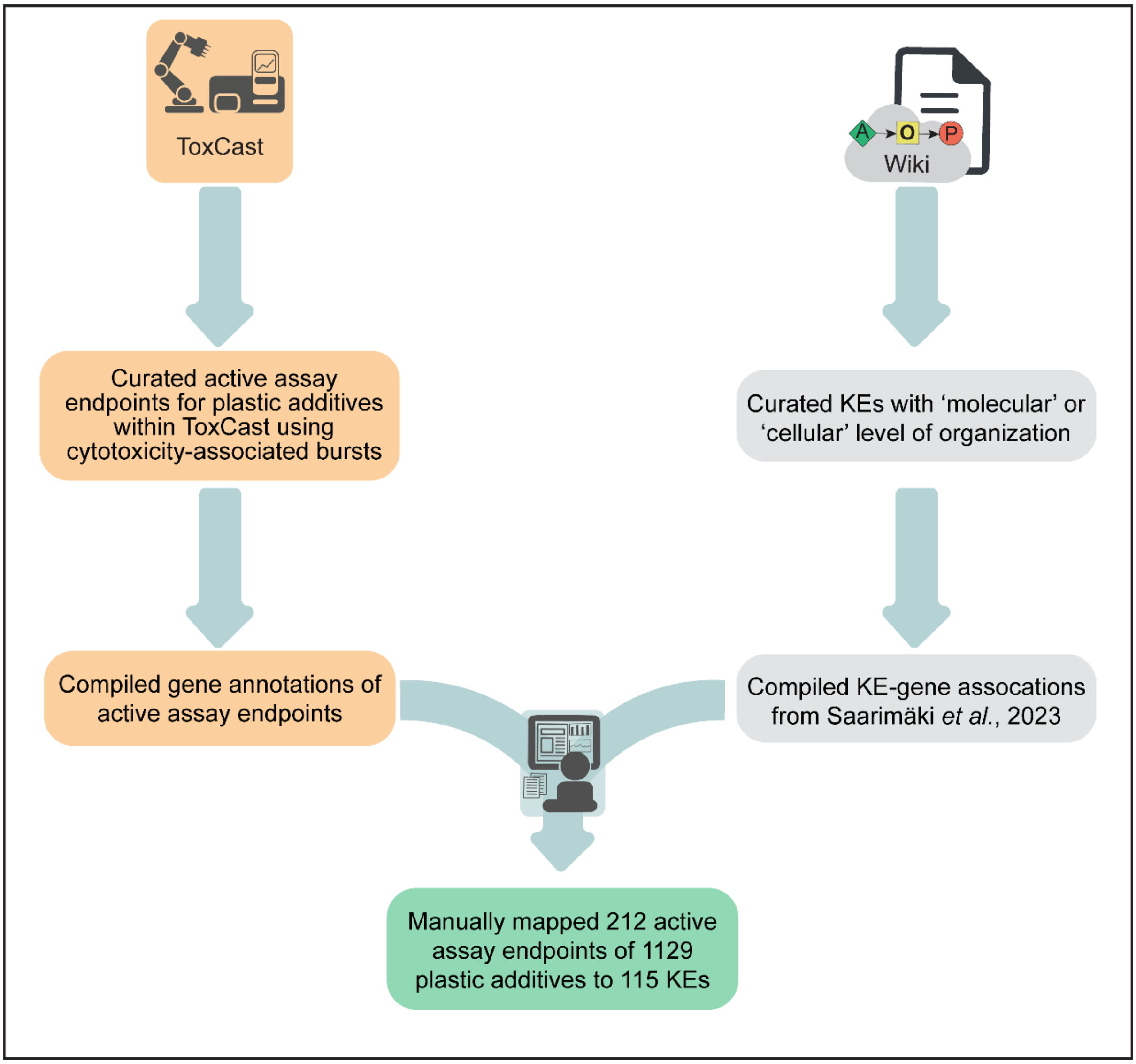


**Figure S2:** Workflow to identify KEs from AOP-Wiki which are mapped to the active assay endpoints of plastic additives within ToxCast.

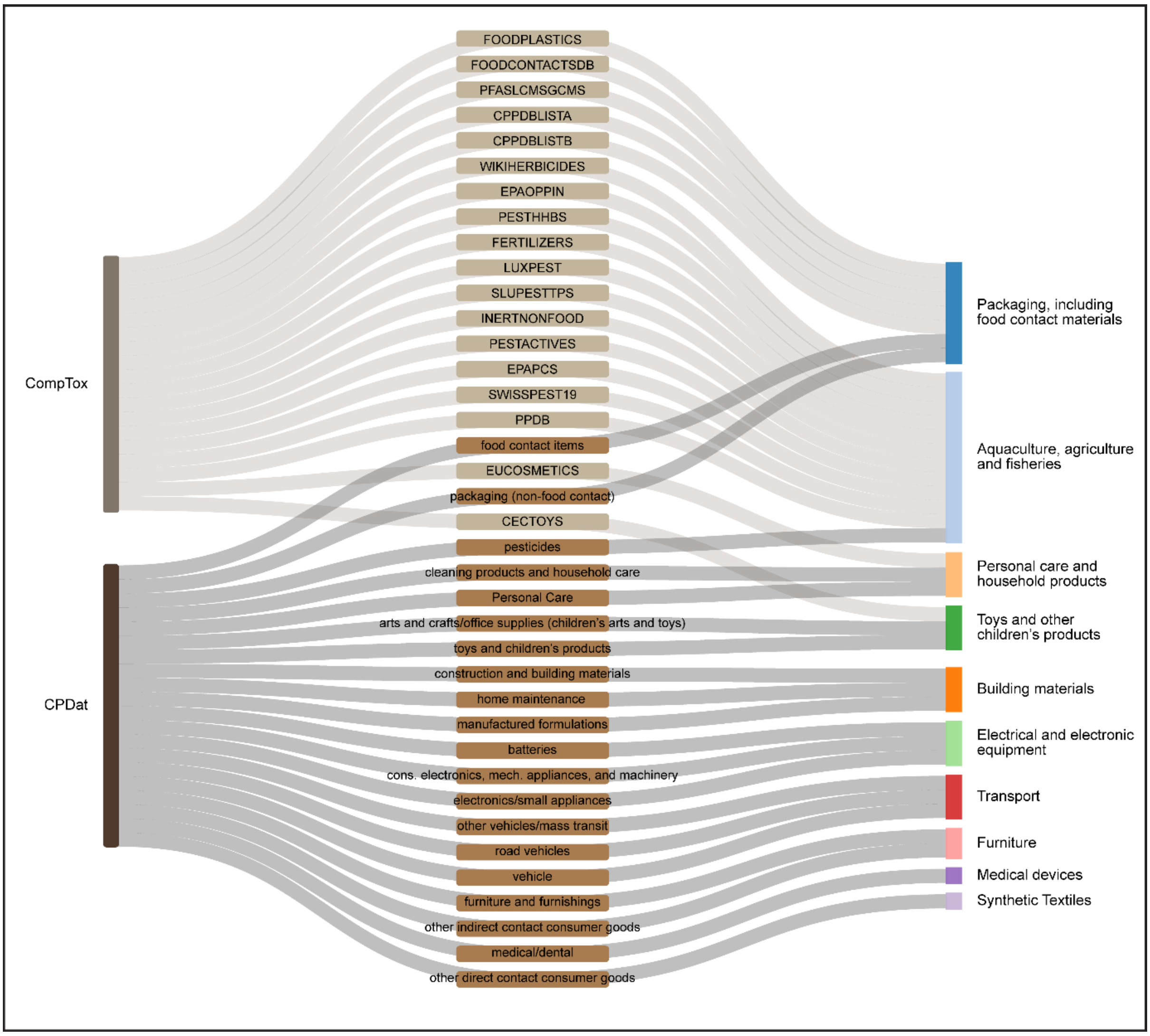


**Figure S3:** Mapping of chemical or category lists from CompTox Chemicals Dashboard and Chemical and Products Database (CPDat) with the 10 priority use sectors of plastic additives.

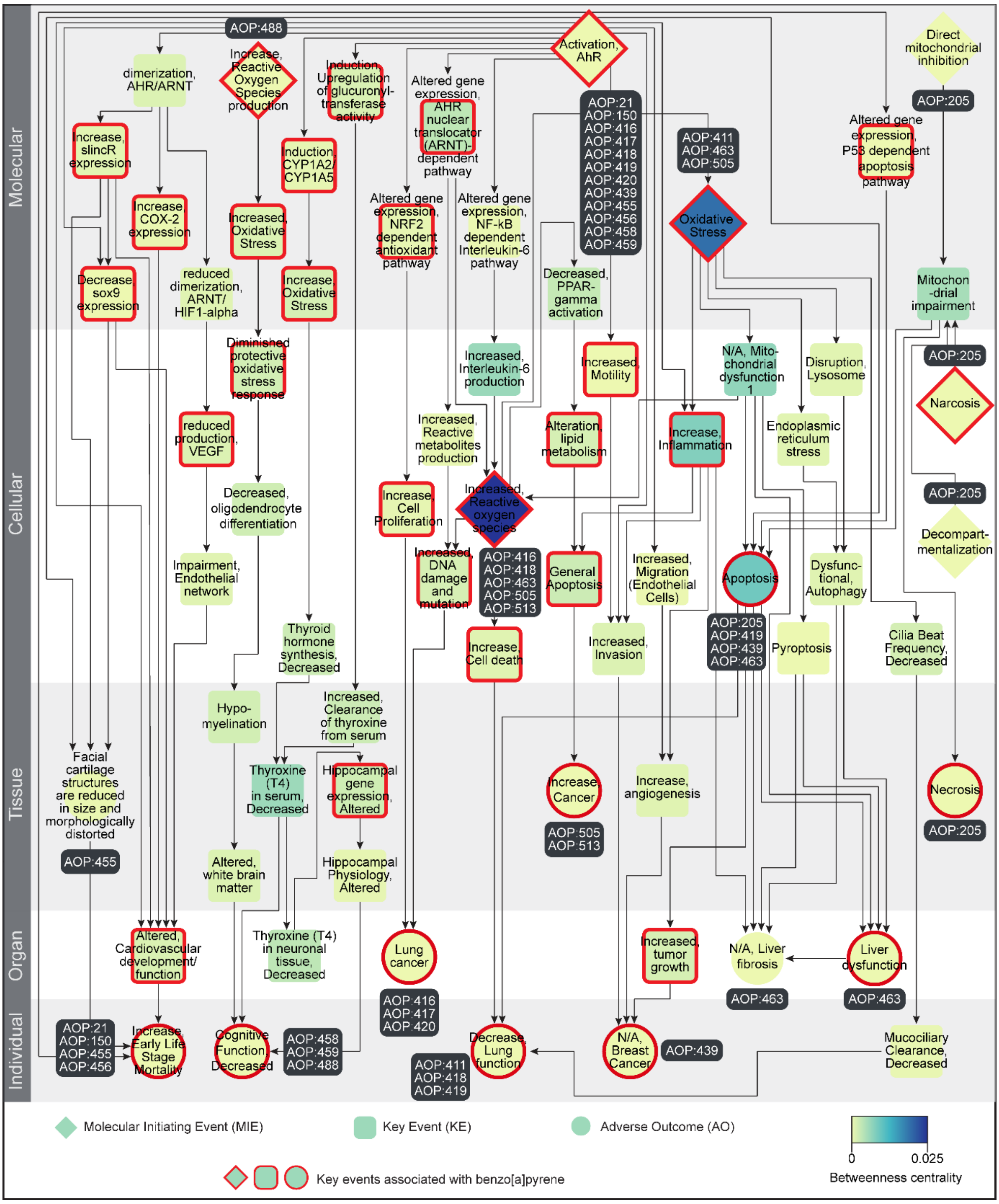


**Figure S4:** Directed network corresponding to the largest connected component (C1) in the B[a]P-AOP network, where the KEs (including MIEs and AOs) are colored based on their betweenness centrality values. The 36 KEs (including MIEs and AOs) associated with B[a]P are marked in 'red'. In this figure, the 66 KEs are arranged vertically according to their level of biological organization.

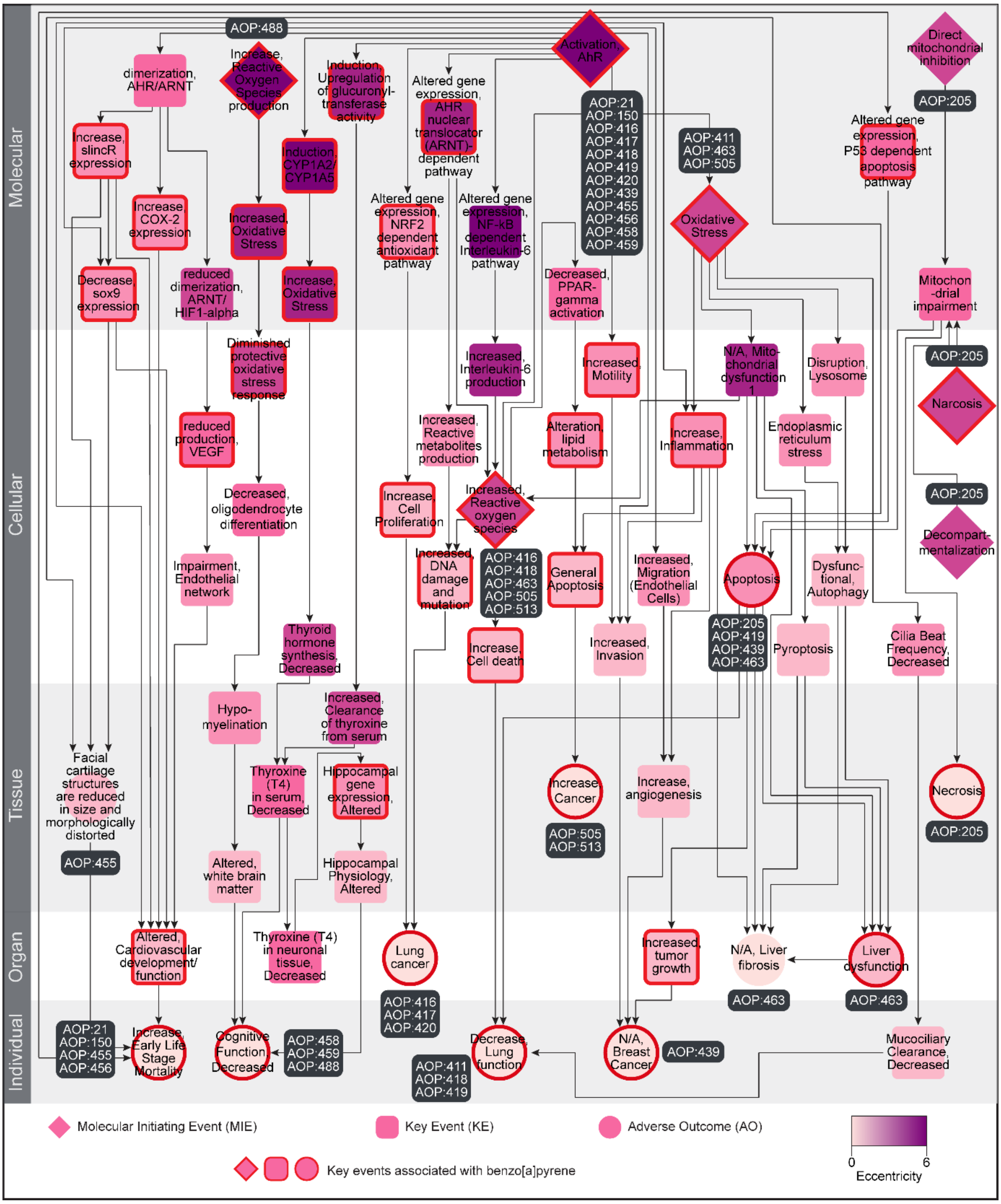


**Figure S5:** Directed network corresponding to the largest connected component (C1) in the B[a]P-AOP network, where the KEs (including MIEs and AOs) are colored based on their eccentricity values. The 36 KEs (including MIEs and AOs) associated with B[a]P are marked in 'red'. In this figure, the 66 KEs are arranged vertically according to their level of biological organization.
